# Chemosymbiotic bivalves contribute to the nitrogen budget of seagrass ecosystems

**DOI:** 10.1101/537746

**Authors:** Ulisse Cardini, Marco Bartoli, Raymond Lee, Sebastian Lücker, Maria Mooshammer, Julia Polzin, Miriam Weber, Jillian M. Petersen

## Abstract

In many seagrass sediments, lucinid bivalves and their sulfur-oxidizing symbionts are thought to underpin key ecosystem functions, but little is known about their role in nutrient cycles, particularly nitrogen. We used natural stable isotopes, elemental analyses, and stable isotope probing to study the ecological stoichiometry of a lucinid symbiosis in spring and fall. Chemoautotrophy appeared to dominate in fall, when chemoautotrophic carbon fixation rates were up to one order of magnitude higher as compared to the spring, suggesting a flexible nutritional mutualism. In fall, an isotope pool dilution experiment revealed carbon limitation of the symbiosis and ammonium excretion rates up to 10-fold higher compared to fluxes reported for non-symbiotic marine bivalves. These results provide evidence that lucinid bivalves can contribute substantial amounts of ammonium to the ecosystem. Given the preference of seagrasses for this nitrogen source, lucinid bivalves’ contribution may boost productivity of these important blue carbon ecosystems.

## INTRODUCTION

Shallow-water chemosynthetic symbioses are widespread where decomposition of organic matter produces sulfide [1]. However, their relevance for ecosystem functioning has received limited attention due to the assumption that chemosynthesis plays a minor role in shallow-water ecosystems. Recent studies are challenging this assumption [2–4]. In seagrass sediments, chemosymbiotic bivalves of the family Lucinidae consume sulfide, allowing more plant growth while relying on the seagrass to stimulate sulfide production by free-living sulfate-reducing microorganisms [3]. Still, we know little about nutrient cycling in lucinid bivalves at both the organismal and the ecosystem scale. Most studies to date have focused on carbon (C) fixation by the symbionts and transfer to the host [5, 6] or on the additional contribution of filter feeding to host nutrition [7]. Nitrogen (N) metabolism has received far less attention until recently, when dinitrogen (N_2_) fixation by chemosynthetic symbionts was shown to be possible in two lucinid species [8, 9]. Concurrently, chemosynthetic symbioses can, to varying degrees, gain their N from ammonium (NH_4_^+^), nitrate, or dissolved free amino acids in their environment [10–12], with the symbionts being able to recycle N waste compounds within the symbiosis [13]. Surprisingly, although these studies attest to the expanded N metabolic versatility of chemosynthetic symbioses, the significance of lucinid bivalves in contributing to their ecosystem N budget has been largely overlooked.

## METHODS

We studied a lucinid bivalve (*Loripes orbiculatus*) in the seagrass (*Posidonia oceanica*) sediments of Elba Island (Italy) during two field expeditions in April (spring) and October (fall) 2016. We analyzed porewater inorganic nutrient concentrations (dissolved inorganic nitrogen – DIN and dissolved inorganic phosphorus – DIP). Stable isotope probing with ^13^C-NaHCO_3_^−^ and ^15^N-N_2_ was used to quantify C and N_2_ fixation by the chemosynthetic symbionts. An isotope pool dilution experiment with ^15^N-NH_4_Cl, was conducted in October to quantify gross and net NH_4_^+^ fluxes by the bivalve symbiosis. Finally, elemental and natural stable isotope analyses were conducted to study the stoichiometric and isotopic niche of host and symbiont under the two contrasting seasons. For details see the Supplementary Methods.

## RESULTS AND DISCUSSION

C fixation by the symbionts was roughly 10-fold higher in October compared to April (p < 0.001; Fig. 1A). N_2_ fixation, measured for the first time here in a chemosynthetic symbiosis using the ^15^N-N_2_ method, also increased in October, although not significantly (Fig. 1B). The biogeochemistry of *P. oceanica* sediments is highly influenced by the seagrass seasonal growth, leaf burial and decay by microorganisms. *P. oceanica* growth shows a late spring maximum and a fall minimum [14], when most nutrients within the sediment are consumed and sulfide accumulates [15]. Our porewater profiles confirm this pattern, with higher DIN concentrations and DIN:DIP ratios in April compared to October (p < 0.01; Fig. S1). *L. orbiculatus* is able to supplement its diet with filter feeding on a seasonal basis [7]. Here we show that not only the host, but also the chemoautotrophic symbionts may modulate their metabolic activities according to the availability of external (or recycled) resources.

**Figure 1.**
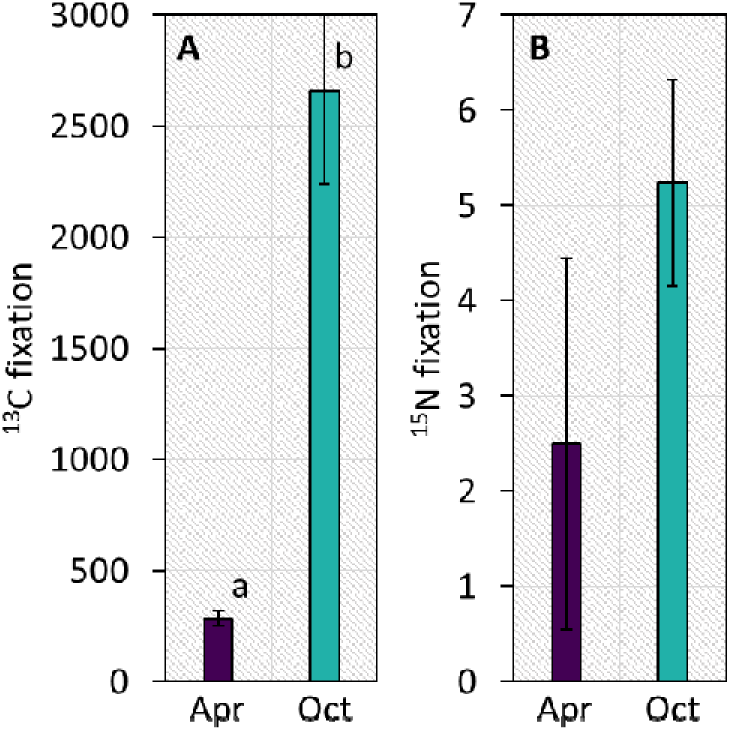
A) Carbon and B) dinitrogen fixation by the microbial symbionts (nmol C (or N) g gill tissue^−1^ h^−1^ ± SE, n = 5). Sampling points are color-coded in purple (April) and cyan (October). Different lowercase letters indicate significant differences.

The increased C fixation rates drove the C:N ratio of the symbionts higher, but not of the host (p < 0.001; Fig. 2A), attesting to the stoichiometric flexibility of the autotrophic partner in the symbiosis and the homeostasis of the heterotrophic host [16]. However, the distribution of bi-variate Bayesian ellipses shows that the natural isotopic niche of the sulfide-oxidizing symbionts was significantly larger in the samples collected in October (Fig. S2), which may indicate a history of mixotrophic metabolism of the endosymbionts, consistent with the presence of a complete tricarboxylic acid (TCA) cycle and transporters for uptake of organic compounds in their genome [8].

**Figure 2.**
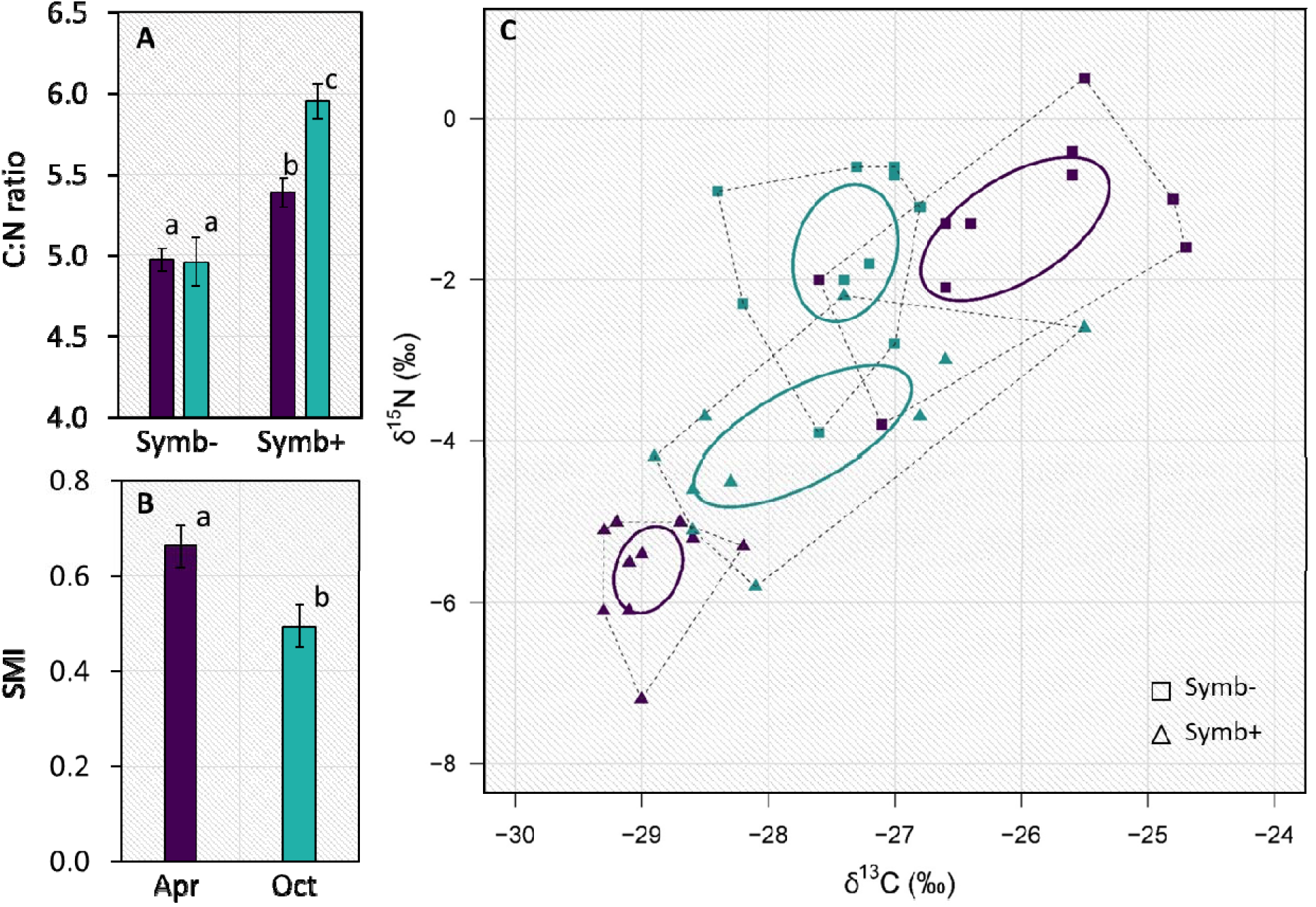
A) C:N ratio (± SE, n = 10) of symbiont-free (Symb-) and symbiont-hosting (Symb+) animal tissues; B) Symbiotic tissue Mass Index – SMI (mg mm^−1^ ± SE, n = 10; see Supplementary Methods for details on how this index was calculated); C) Biplot of the natural abundance of ^13^C and ^15^N isotopes showing the total amount of niche space occupied (Total Area, dashed polygons) and the isotopic niche breadth (Standard Ellipse Area, solid ellipses) of symbiont-free (squares) and symbiont-hosting (triangles) animal tissues. Sampling points are color-coded in purple (April) and cyan (October). Different lowercase letters indicate significant differences.

The proportion of symbiont-hosting gill biomass was lower in October (p < 0.05; Fig. 2B). At the same time, there was a strong overlap in C isotopic niche of host and symbionts, while there was a mismatch in April (Fig. 2C). This could be explained by a flexible nutritional mutualism. Under nutrient rich/high productivity conditions in April, when labile organic matter in seagrass sediments is highest [17], the host relies more on mixotrophy through filter feeding. Under nutrient depleted/low productivity conditions in October, the symbiosis shifts towards relying more on the symbionts as a source of energy. Our observation that symbiont C fixation rates were 10 times higher in October compared to April is consistent with this theory. Filter-feeding bivalves that do not host chemosynthetic symbionts enter a ‘dormant’ state in summer, possibly due to food limitation [18 and references therein]. The ability to harvest energy throughout the summer and fall by relying on symbiont primary production when food availability is low would provide lucinid bivalves with a distinct advantage over non-symbiotic filter-feeding bivalves. While more targeted approaches will be needed to conclusively verify this hypothesis, gross NH_4_^+^ production and consumption measured in October using isotope pool dilution (IPD) indicated that the symbiosis was indeed C limited, as bivalves consumed NH_4_^+^ only when exposed to a source of labile organic C (Fig. S3).

The same experiments, using IPD on an invertebrate symbiotic animal for the first time to our knowledge, allowed us to quantify gross and net excretion rates contributed by the symbiosis to its surroundings. Net excretion by the bivalves was approximately 15 µmol NH_4_^+^ g_SFDW_^−1^ h^−1^ (Fig. S3), which is up to 10-fold higher compared to NH_4_^+^ excretion rates reported for other non-symbiotic marine bivalves [19] and testifies to the potential of these chemosynthetic symbioses to underpin ecosystem functioning by nitrogen provisioning.

## CONCLUSIONS

In this study, we show that *L. orbiculatus* likely has a flexible nutritional mutualism, in which host and symbionts cycle between a looser trophic association and a tight chemoautotrophic partnership, changing nutritional strategy according to the environmental conditions. Further, we report that under C-limiting conditions these chemosymbiotic bivalves can excrete substantial amounts of NH_4_^+^ to the environment. In seagrass sediments, lucinids and their endosymbionts are not only relevant for their role in sulfide detoxification [3], but can also provide the plant’s preferred N form [20], thus contributing to the productivity of these important blue carbon ecosystems.

## ACKNOWLEDGEMENTS

The authors thank Margarete Watzka for her assistance with sample preparation for IRMS analyses, and Alexandra Belitz, Nathalie Elisabeth, Christian Lott and the team of the HYDRA Field Station Elba for their assistance during fieldwork and/or lab activities. Augusto Passarelli is kindly acknowledged for his help with the inorganic nutrient analyses. The Petersen laboratory is supported by a Vienna Research Groups for Young Investigators grant from the Vienna Science and Technology Fund (WWTF). U.C. and M.B. are supported by the project INBALANCE (No. 09.3.3-LMT-K-712-01-0069), funded by the Research Council of Lithuania (LMT) under the European Social Funds (ESF) programme.

## AUTHOR CONTRIBUTIONS

U.C. conducted the fieldwork and performed all the experiments, analyzed the data and wrote the manuscript. M.B. conducted ^15^N- NH_4_^+^ measurements at the membrane inlet mass spectrometer, and provided critical input for data interpretation. R.L. conducted bulk stable isotope measurements on bivalve tissues and provided critical input for data interpretation. S.L. provided guidance for GC-MS measurements of ^15^N enrichments in seawater samples. M.M. assisted in designing the IPD experiment and provided critical input for data interpretation. J.P. contributed to the project during fieldwork and sample analyses. M.W. assisted with the organization and conduction of all fieldwork activities. J.M.P. contributed to the design of the research project, data interpretation, and manuscript production.

