## Supplementary information for "Chemosymbiotic bivalves contribute to the nitrogen budget of seagrass ecosystems"

### 1. SUPPLEMENTARY FIGURES

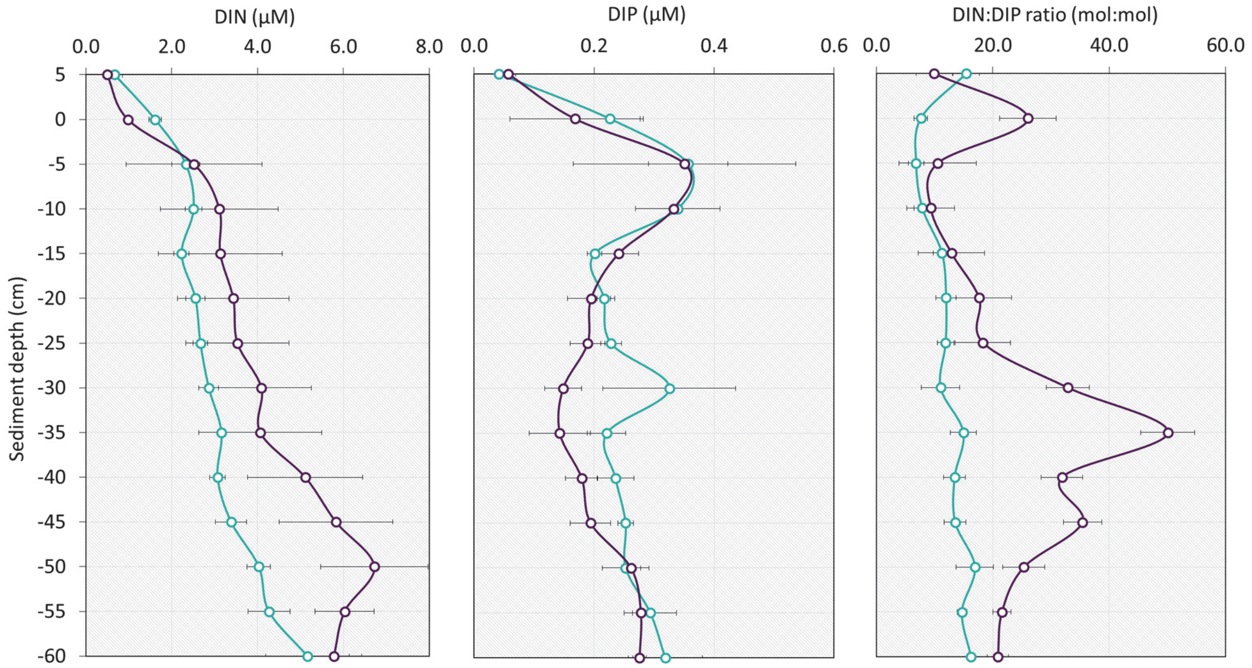


**Figure S1**: Dissolved inorganic nutrients and DIN:DIP ratio in the porewater at the time and place of sampling (± SE, n = 3), color-coded in purple (April) and cyan (October).

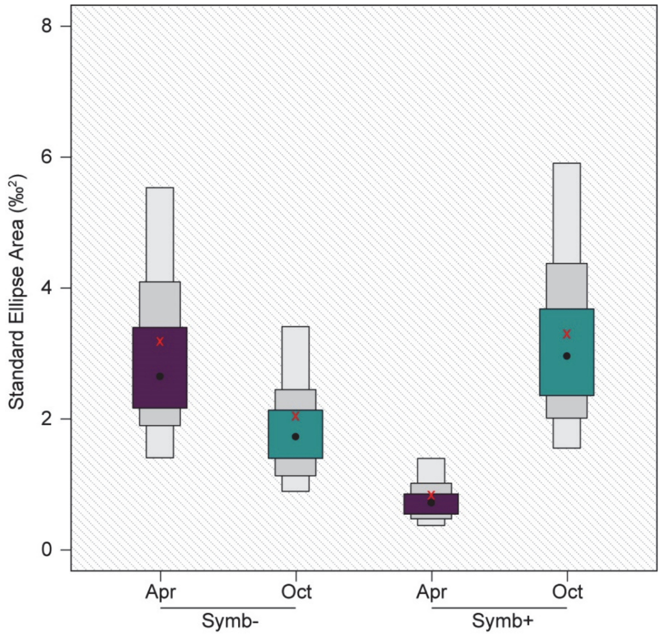


**Figure S2**: Distribution of Bayesian ellipses showing the isotopic niche breadth and its uncertainty for symbiont-free (Symb-) and symbiont-hosting (Symb+) animal tissues, color-coded in purple (April) and cyan (October). Black dots represent the mode while red crosses are population means and the shaded boxes represent the 50%, 75% and 95% credible intervals from dark to light grey.

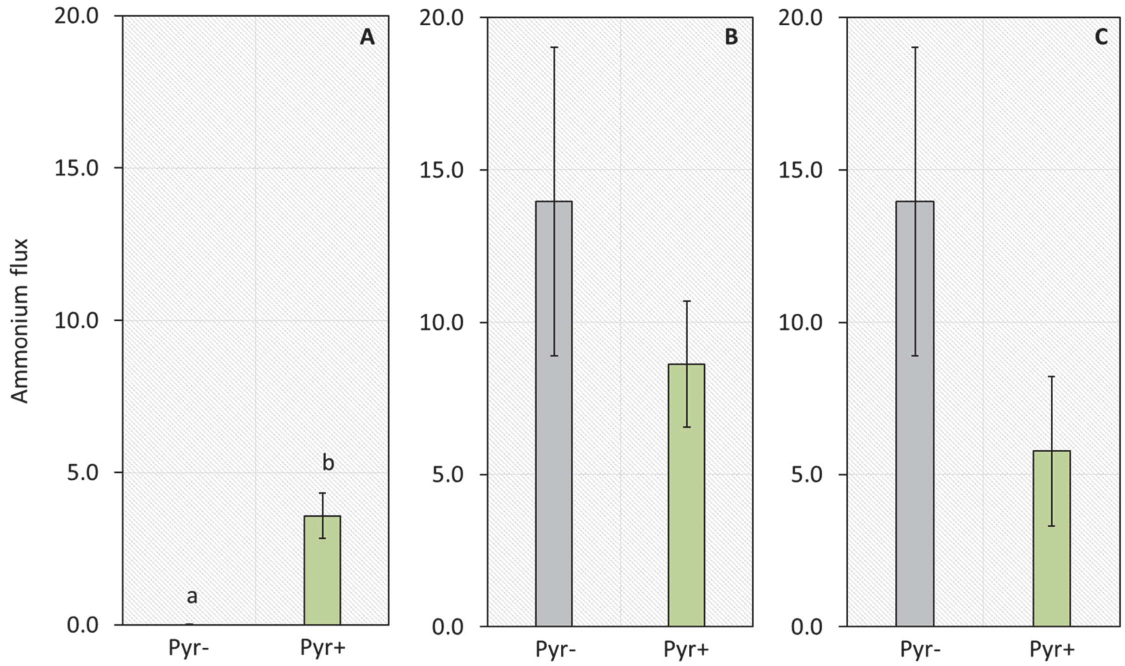


**Figure S3**: Ammonium fluxes (μmol $NH_4^+$ $g^{-1}$ $h^{-1}$ ± SE, n = 5) measured during the isotope pool dilution experiment in October. A) Gross uptake; B) Gross excretion; C) Net excretion. Results for the incubations with natural filtered seawater (Pyr-) are reported in grey, while results of the incubations amended with 10 μM pyruvate (Pyr+) are reported in green. Different lowercase letters indicate significant differences.

### 2. SUPPLEMENTARY METHODS

#### Field collection and porewater nutrients

All the sampling activities and the incubation experiments were conducted during two field expeditions to the Elba Island station of the HYDRA Institute for Marine Sciences in Fetovaia, Livorno (Italy) in April and October 2016. Specimens of *Loripes orbiculatus* were collected by scuba diving in the bay of Fetovaia from sediments adjacent to a *Posidonia oceanica* meadow (42°43'48"N 10°9'23"E) at approximately 7 m depth. The bivalves were transported back to the Station with a good quantity of their surrounding sediment within 30 minutes and placed in the shade at ambient temperature in 30 L aquaria with natural seawater to acclimatize before the incubation experiments.

Porewater was collected by scuba divers at the sampling location in the bay using metered stainless steel lances designed to penetrate down to 60 cm below the sediment surface, with a resolution of 5 cm between collected samples. In each season, three profiles were collected for a total of 42 samples (14 x 3). Porewater samples were filtered on 0.22 μm polycarbonate membrane filters (Merck Millipore), preserved frozen at -20°C and finally analyzed for nitrate, nitrite, ammonium, and orthophosphate concentrations on a Continuous Flow Autoanalyzer (Flowsys, Systea s.p.a.) at the Stazione Zoologica Anton Dohrn, Napoli, Italy.

#### $^{13}C$-$HCO_3^-$ and $^{15}N$-$N_2$ isotope probing experiments and stable isotope analysis

Isotope probing experiments were conducted with lucinid bivalves to quantify C and $N_2$ fixation by the chemosynthetic symbionts during the two field campaigns. All material used for the incubation experiments was soaked in 2 M HCl overnight and rinsed with MilliQ water before use. $^{15}N$- and $^{13}C$-enriched seawater was prepared before the incubation. $^{13}C-NaHCO_3^-$ (98 atom% 13C, Sigma Aldrich) was dissolved in 0.2-µm-filtered seawater to reach a $H^{13}CO_3^-$ concentration of 2 mM (final $^{13}$C-atom% = 51.6%). $^{13}$C-enriched seawater (or unamended natural abundance seawater, for the controls) was transferred to 250 ml serum bottles, crimp-sealed gas-tight with a hollow needle in the septum to prevent air bubbles forming. Thereafter, 10 ml of $^{15}N-N_2$ gas (99 atom% $^{15}$N, Cambridge Isotope Laboratories, lot number: 01/071401) (or of air, for the controls) were injected in the serum bottles with a hollow needle in the septum to allow replacement of the liquid. The procedure was repeated again with additional 20 ml of gas without the second needle in the septum, to create over-pressure within the bottles and aid dissolution of the gas [1, 2]. This modified procedure, without further dilution, resulted in a final $^{15}$N-atom% of 47.4% (for details of the measurement see the following). The serum bottles were shaken for 2 minutes and stored in the dark, upside down, until use. The $^{15}N-N_2$ gas was checked for the presence of $^{15}$N-nitrate or $^{15}$N-ammonium at the Max Planck Institute for Marine Microbiology, Bremen, Germany, and was found to be free of contamination [3].

25 ml serum bottles were prepared with a layer of 1.5 cm of acid-washed glass beads (425-600 µm, Sigma Aldrich) at the bottom to simulate the sediment in which the bivalve burrows. Each serum bottle was then filled with the respective treatment seawater (control or enriched), gently transferring the enriched seawater to minimize loss of $^{15}N-N_2$ gas. Bivalves of ca. 1 cm shell height (adult size) were subsequently added and the bottles crimp-sealed gas tight avoiding the formation of bubbles. Five bivalves were incubated either in $^{15}$N- and $^{13}$C-enriched seawater or in control seawater.

Initial samples (n = 3) were collected from each 250 ml serum bottle in 12 ml exetainers (Labco Limited), preserved with 50 µl of 0.25M $HgCl_2$ solution and stored upside down in the dark until analysis. The isotopic composition of the $N_2$ gas in the enriched seawater was analyzed after headspace equilibration using gas chromatography (Agilent 6890 equipped with a Porapak Q column at 80 °C and a TCD detector at 300 °C; Agilent Technologies, Santa Clara, CA, USA) combined with mass spectrometry (Agilent 5975c, quadruple inert MS) as in [4]. After 24 h, the incubation bottles were opened, the bivalves collected, measured for their shell length and dissected. Symbiont-bearing (gill) tissue and non-symbiotic (host) tissue were separated and stored at -20°C. Ten lucinid specimens were dissected and preserved without any incubation both in April and in October to determine the natural $^{13}C/^{12}C$ and $^{15}N/^{14}N$ ratios of the gill and host tissues. Frozen tissues were freeze-dried for 48 h before being ground to fine powder and weighed into tin capsules that were crimped manually. Samples were analyzed for carbon and nitrogen elemental composition (%) and isotope ratios ($\delta^{13}C$ and $\delta^{15}N$) by continuous flow isotope ratio mass spectrometry (IRMS) using a Costech elemental analyzer interfaced with a GV Instruments Isoprime IRMS. Measures of $\delta^{13}C$ and $\delta^{15}N$ of the samples were compared against reference materials with a precision and accuracy of 0.3 and 0.5 ‰, respectively. $^{15}N_2$ and $^{13}$C incorporation rates were calculated following [2, 5], with the equation:

$$\frac{\text{gill } ^{15}\text{N (or } ^{13}\text{C) atom\% excess}}{\text{medium } ^{15}\text{N (or } ^{13}\text{C) atom\% excess}} \times \frac{\mu\text{g gill N (or C)}}{\text{g gill ind}^{-1}} \times \frac{1}{\Delta t}$$

and expressed as nmol N (or C) g gill tissue$^{-1}$ h$^{-1}$ ± SE.

**$^{15}N$-$NH_4Cl$ isotope pool dilution experiment**

In October, we conducted an isotope pool dilution experiment to quantify gross and net $NH_4^+$ fluxes by the bivalve symbiosis. This technique is based on labeling the ammonium pool by adding $^{15}N$-labelled ammonium. The quantification of the decrease in the isotopic label and the change in concentrations over time allows to calculate the gross production (i.e., mineralization) and immobilization rates. $^{15}N$-$NH_4Cl$ (98 atom% $^{15}N$, Sigma Aldrich) was added to 0.2-µm-filtered seawater to reach a $^{15}N$-$NH_4^+$ concentration of 0.4 µM. Thereafter, the incubation experiment consisted of 3 different sets of 25 ml serum bottles (n = 5 per treatment/control) prepared with a layer of 1.5 cm of acid-washed glass beads as described above. The control consisted of serum bottles with $^{15}N$-$NH_4^+$-enriched seawater but without bivalves. In the second set, the bivalves were added. The third set was additionally amended with sodium pyruvate ($C_3H_3NaO_3$, Sigma Aldrich, final concentration: 10 µM) to provide the bivalves with a source of labile organic C. Initial samples were taken from the stock solution used to fill each set of bottles, and after 6 h of incubation from each serum bottle. The samples were collected in 12 ml exetainers, preserved with 50 µl of 0.25M $HgCl_2$ solution and stored upside down in the dark until analysis. Each sample was analyzed for total $NH_4^+$ concentrations (with a standard spectrophotometric technique) and for $^{15}N$:$^{14}N$ ratios of $NH_4^+$. To this purpose, samples in exetainers were degassed with helium, treated with a hypobromite-iodine solution to oxidize $NH_4^+$ to $N_2$, and analyzed on a membrane-inlet mass spectrometer (Bay Instrument). Gross immobilization (i, uptake) and mineralization (m, excretion) rates by the bivalve symbiosis were calculated using the equations by Kirkham & Bartholomew [6], with net excretion = m-i. For the control samples without bivalves, the immobilization and mineralization rates were not different from zero, because no dilution of the isotopic label over time occurred, due to lack of activity.

**Elemental and natural stable isotope analyses**

C% and N% data obtained after analysis of non-incubated tissue samples at the EA-IRMS was used to calculate C:N ratios of the gill and host tissues. The Symbiotic tissue Mass Index was determined as the ratio between the gill tissue dry weight of each bivalve specimen (mg) and its shell length (mm). Individual $\delta^{13}C$ and $\delta^{15}N$ values of symbiont-bearing and non-symbiotic tissues were analyzed using a Stable Isotope Bayesian Ellipses in R (SIBER) model [7, 8] to compare isotopic niche spaces of symbionts and host in April and October. The SIBER Bayesian model generates a bi-plot with standard ellipses that contain approximately 40% of the data and are insensitive to sample size and robust measures of isotopic niche width useful in comparative studies between groups [7]. The distribution of Bayesian ellipses was subsequently plotted for each group as a density plot.

**Statistical analyses**

Differences in each parameter were assessed using univariate distance-based permutational nonparametric analyses of variance (PERMANOVA) [9]. Environmental variables (DIN, DIP, DIN:DIP) were normalized and analyzed for differences between seasons based on Euclidean distances using type I (sequential) sum of squares with 9999 unrestricted permutations of raw data. All other variables were square root transformed, and analyses were based on Bray Curtis similarities using type III (partial) sum of squares with 9999 unrestricted permutations of raw data. Uptake and excretion rates were tested for differences between treatments (with or without pyruvate amendment), while the Symbiotic tissue Mass Index, C fixation and N fixation rates for differences between seasons. A fully crossed design with two fixed factors (seasons, tissues) was used to test for differences in C:N ratios. Pair-wise tests were carried out if significant differences occurred ($P<0.05$). PERMANOVA tests were performed in the software PRIMER 6+ (PRIMER-E Ltd, Plymouth, UK).
